# Modular Engineering of Thermo-Responsive Allosteric Proteins

**DOI:** 10.1101/2025.05.02.651844

**Authors:** Kira H. Hoffmann, Ann-Sophie Kroell, Nikolas A. Motzkus, Nina Lemmen, Nele Happ, Benedict Wolf, Anna von Bachmann, Nicholas Southern, Felicitas Vogd, Sabine Aschenbrenner, Dominik Niopek, Jan Mathony

## Abstract

Thermogenetics enables non-invasive spatiotemporal control over protein activity in living cells and tissues, yet its applications have largely been restricted to transcriptional regulation and membrane recruitment. Here, we present a generalizable strategy for engineering thermosensitive allosteric proteins through the insertion of optimized *Avena sativa* LOV2 domain variants. Applying this approach to a diverse set of structurally and functionally unrelated proteins in *Escherichia coli*, we generated potent, thermo-switchable chimeric variants that can be tightly controlled within narrow temperature ranges (37-41°C). Extending this strategy to mammalian systems, we engineered the first CRISPR-Cas genome editors directly modulated by subtle temperature changes within the physiological range. Finally, we showcase the incorporation of a chemoreceptor domain as an alternative thermosensing module, suggesting thermo-sensitivity to be a widespread feature in receptor domains. This work expands the toolkit of thermogenetics, providing a blueprint for temperature-dependent control of virtually any protein of interest.

## Introduction

Engineered switchable proteins are widely applied in basic research and biotechnology. They are typically based on naturally occurring proteins that are modified to be activated or deactivated by exogenous stimuli, most commonly light or chemicals, allowing researchers to remotely control various cellular processes. With respect to future biomedical applications, temperature represents a particularly attractive physical cue due to the increased tissue penetration compared to light and the superior spatiotemporal precision compared to chemicals^1–3^. Efficient approaches for engineering thermo-responsive proteins are lacking, however, and the few existing approaches are mostly limited to the control of gene expression^4–6^. Recent work employed the *Bc*LOV4 domain for temperature-dependent recruitment of proteins to the plasma membrane^7,8^. While this approach is generally compatible with various different effector proteins, it relies on a large protein domain (*Bc*LOV4 is ∼600 aa) and is restricted to effectors, the activity of which depends on plasma membrane localization.

To address these key limitations, we aimed to develop a generalizable approach for thermogenetic control via modular engineering of temperature-dependent protein allostery. By inserting the *Avena sativa* LOV2 domain (*As*LOV2) from phototropin-1 and improved mutants thereof into effector proteins, we demonstrate the modular engineering of thermo-responsive hybrids. Starting with the bacterial transcription factor AraC as a proof-of-concept, we achieved potent and tunable activity switching in response to small temperature shifts between 37°C to 41°C. We successfully transferred this concept to several additional proteins, including different CRISPR-Cas9 gene editors and transactivators, and showcase the broad applicability of the concept in bacteria and in mammalian cells. Finally, we extend the method to a second receptor using a glucocorticoid ligand-binding domain and demonstrate the dual regulation of genome editing by the FDA-approved drug cortisol and temperature. Our work overcomes a major bottleneck in thermogenetics by introducing the first general approach for engineering thermogenetic switches that can be controlled within the narrow human physiological temperature range and are not limited to specific applications or protein classes.

## Results

### Engineering a thermo-switchable transcription factor by *As*LOV2 insertion into AraC

An ideal module would comprise a compact protein domain that exerts a large conformational rearrangement in response to small changes in temperature. Insertion of such a domain into an effector protein of choice could thus couple the activity of the effector to the conformational state of the thermosensor. Intrigued by the relatively low stability of many LOV domains^7,9,10^, we decided to test the widely applied *As*LOV2 for its response to temperature changes. To this end, we used an optogenetic variant of the transcription factor AraC carrying an *As*LOV2 insertion after residue S170, which we had previously established for co-regulating transgene expression by light and arabinose in *E. coli*^11^ (Fig. 1a). At 37°C, both wild-type AraC and the AraC-LOV domain hybrid mediated strong expression of a pBAD-driven, monomeric red fluorescent protein (RFP) (Supplementary Fig. 1). At 41°C, however, strongly reduced RFP expression was consistently observed for AraC-LOV, whereas the wild-type AraC control remained unaffected, indicating a potent thermo-response of AraC-LOV (Fig. 2 a,b and Supplementary Fig. 1). Importantly, cell growth was only slightly affected by the elevated temperature (Supplementary Fig. 1). Furthermore, mutation of the *As*LOV2 C450 residue, which is central to the native *As*LOV2 photocycle, to alanine did not affect the temperature switching of the AraC-LOV hybrid and was included in all downstream constructs to avoid cross-reaction with ambient light.

**Figure 1.**
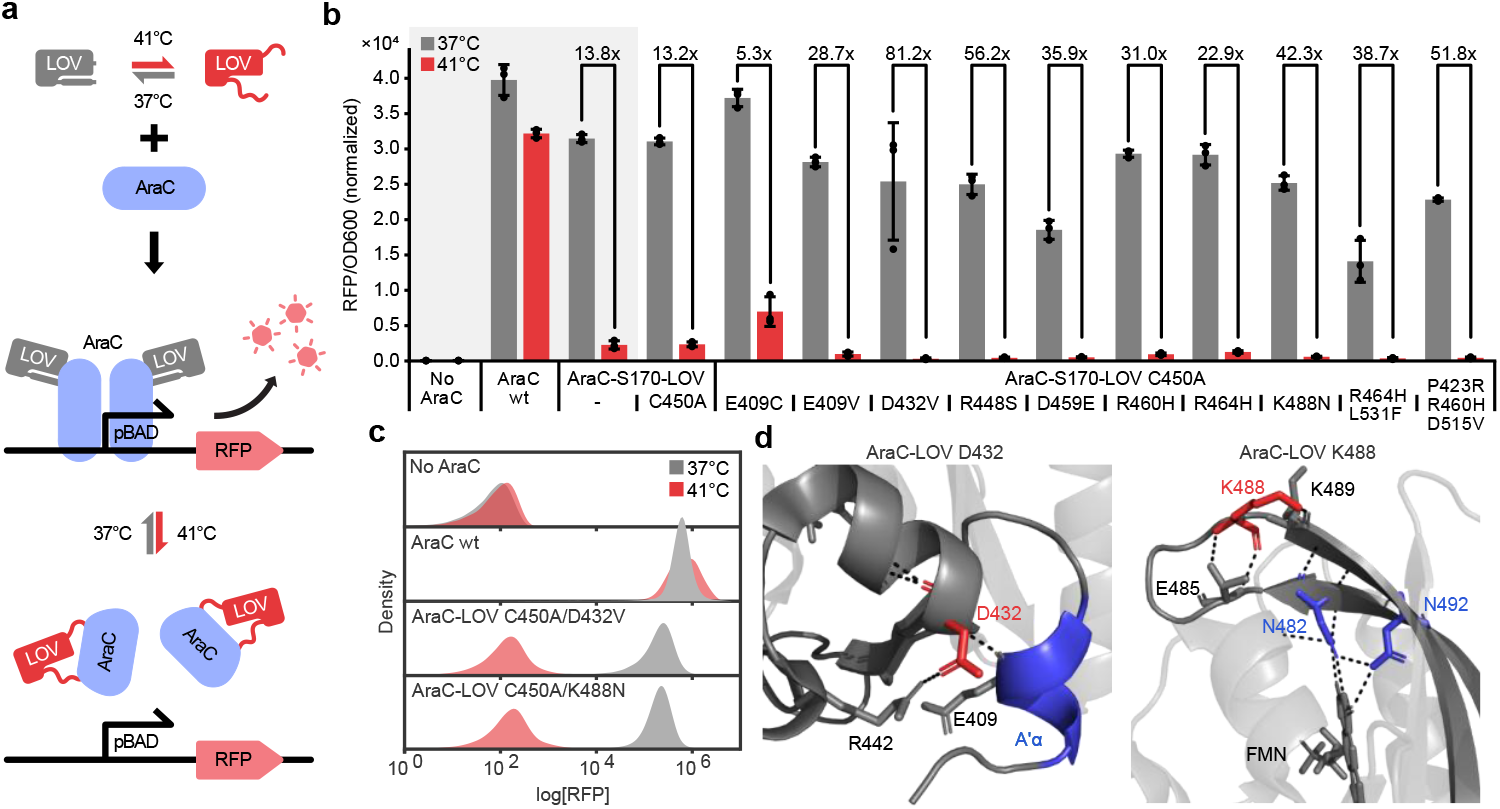
Engineering a thermo-switchable *E. coli* transcription factor by *As*LOV2 domain insertion. **a**, Assay schematics. **b**,**c**, *E. coli* transformed with plasmids encoding the indicated AraC-LOV variants and an mRFP reporter were incubated at 37°C or 41°C followed by measurements of reporter fluorescence (RFP) and OD600 in a plate reader (**b**), or flow cytometry (**c**). **d**, Close-ups of the *As*LOV2 crystal structure. Mutated residues are marked in red and functionally critical elements are marked in blue. Hydrogen bonds are indicated. PDB-ID: 2V0U. **b** Data represent the mean +/-SD, n = 3 independent experiments.

**Figure 2.**
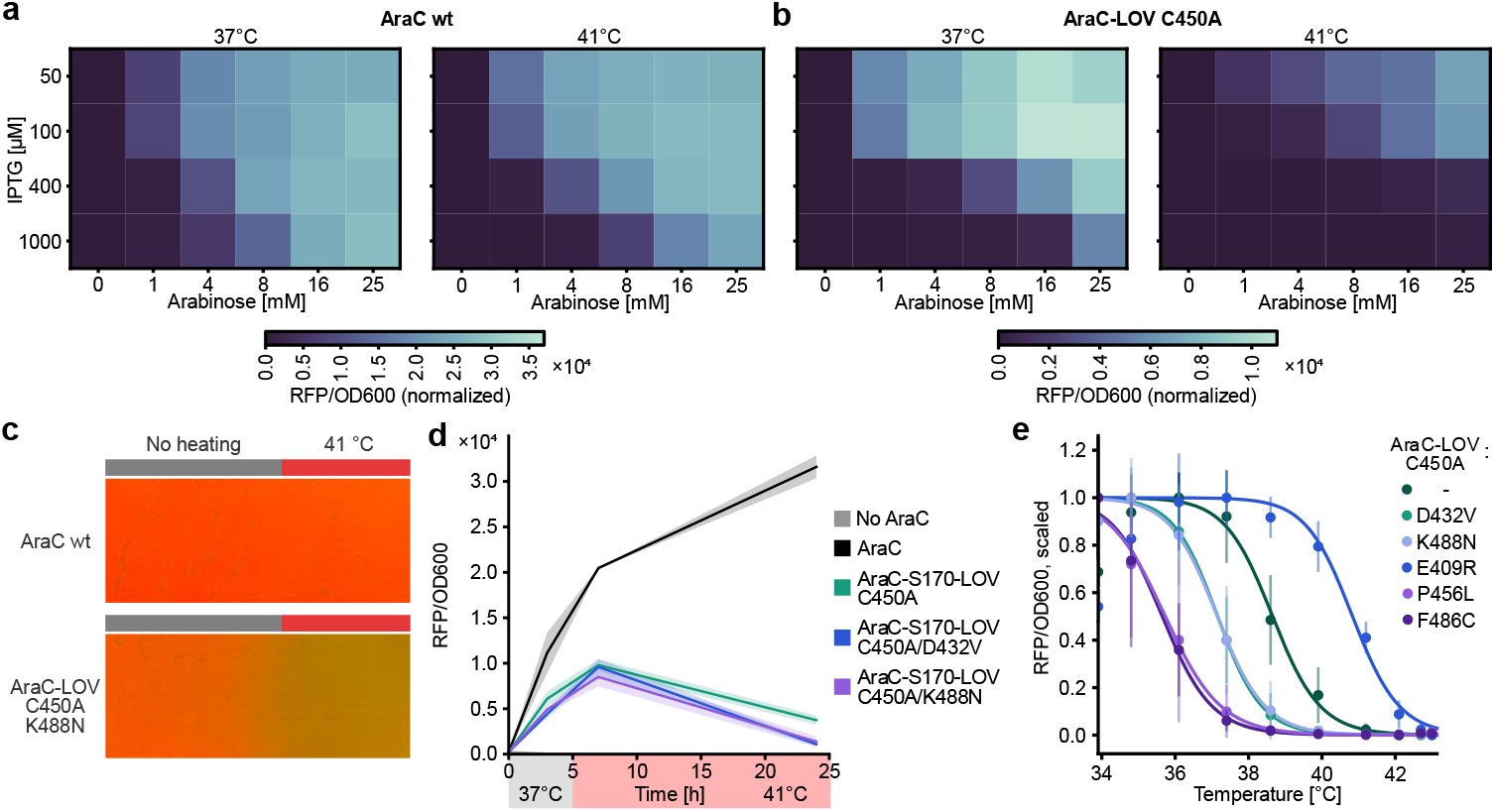
Thermosensitive AraC-LOV hybrids are precisely tunable and enable spatiotemporal control of gene expression. **a, b**, *E. coli* carrying a pBAD-mRFP reporter and AraC wt (**a**) or AraC-LOV C450A (**b**) were incubated for 16 h at 37°C or 41°C in the presence of inducers at the indicated concentration. Expression of the AraC variants is IPTG-inducible, while AraC activity is arabinose-dependent. RFP fluorescence and OD600 were evaluated in a plate reader. **c**, Photographs of spatial gene expression control. AraC-(LOV) reporter strains were plated on agar and one half of the tray was heated while the other half was kept at room temperature. **d**, *E. coli* containing a pBad-mRFP reporter and expressing the indicated AraC variant or a dummy control protein were incubated at 37°C. After 5 hours the temperature was increased to 41°C followed by another 19 hours of incubation. Culture density (OD600) and RFP expression (normalized to OD) were periodically assessed in a plate reader. **e**, Temperature response profiles of AraC-LOV variants. Samples were prepared as in **a** but incubated at different temperatures. Data was min-max normalized for each sample (see Supplementary Fig. 6). Data represent the mean +/-SD, n = 3 independent experiments. wt, wild-type.

Next, to optimize the dynamic range of thermal control, we performed error-prone PCR on LOV-C450A and screened the resulting library of *E. coli* colonies (complexity ∼5×10E6) for temperature-dependent reporter expression on agar plates (see Online Methods and Supplementary Fig. 2). We identified several *As*LOV2 point mutants that significantly improved the dynamic range of the system by reducing reporter expression in the 41°C off state, as indicated by plate reader measurements and flow cytometry (Fig. 1b,c and Supplementary Fig. 3). The two lead candidates, D432V and K488N, showed 81- and 42-fold changes in reporter expression upon temperature induction, respectively. Of note, these mutations are likely to have destabilizing effects: D432 forms a hydrogen bond with E409 in the mechanistically important A’α helix (Fig. 1d), while K488 stabilizes a loop connecting two β-sheets, both in direct contact with the flavin mononucleotide (FMN) chromophore (Fig. 1d).

We continued with a detailed characterization of our thermogenetic system. Inducer dose escalation experiments revealed that the thermal response of AraC-LOV C450A is expression-dependent and can be tuned over a wide range of inducer concentrations (Fig. 2a,b and Supplementary Fig. 4). Next, focusing on our lead candidates, we showcased efficient spatial control when *E. coli* expressing AraC-LOV-C450A/K488N were grown on agar plates and a heat gradient was applied (Fig. 2c). Moreover, we demonstrated reversible temperature-control via timed incubation of liquid cultures at different temperatures for both the D432V and K488N mutant (Fig. 2d and Supplementary Fig. 5). Subsequent experiments, in which identical samples were incubated at several temperatures between 34-43°C in parallel, revealed a sharp decrease in transcriptional activation within a range of only 3-4°C for the improved AraC-LOV hybrids (Fig. 2e). Using our screening pipeline, we further identified variants with shifted transition temperatures in the range of 34-37°C, thus even allowing cold-induced protein activation at temperatures below 37°C (Fig. 2e and Supplementary Fig. 6).

### Allosteric thermo-control is applicable to diverse bacterial protein classes

Next, to test the modularity of our thermogenetics approach, we transferred the concept to different effector proteins, starting with a chloramphenicol acetyltransferase (CAT, Fig. 3a). Building on an existing optogenetic *As*LOV2 insertion variant at K136 of the antibiotic resistance enzyme^12^, we evaluated effects of different flexible and rigid interdomain linkers around the *As*LOV2 insertion site, and assessed these variants using cell growth assays in the presence of chloramphenicol as a readout (Fig. 3b and Supplementary Fig. 7). Cultures expressing CAT-LOV hybrids reliably reached stationary phase at 37°C, albeit at a slower growth rate than the wild-type CAT control (Supplementary Fig. 7a). Excitingly, for the CAT-LOV-C450A hybrid with LOV-flanking GP linkers and some *As*LOV2 mutants thereof, antibiotic resistance was effectively switched off at 41°C resulting in cell death, as indicated by OD600 measurements and colony titers of serial dilutions (Fig. 1c-e and Supplementary Fig. 7b).

**Figure 3.**
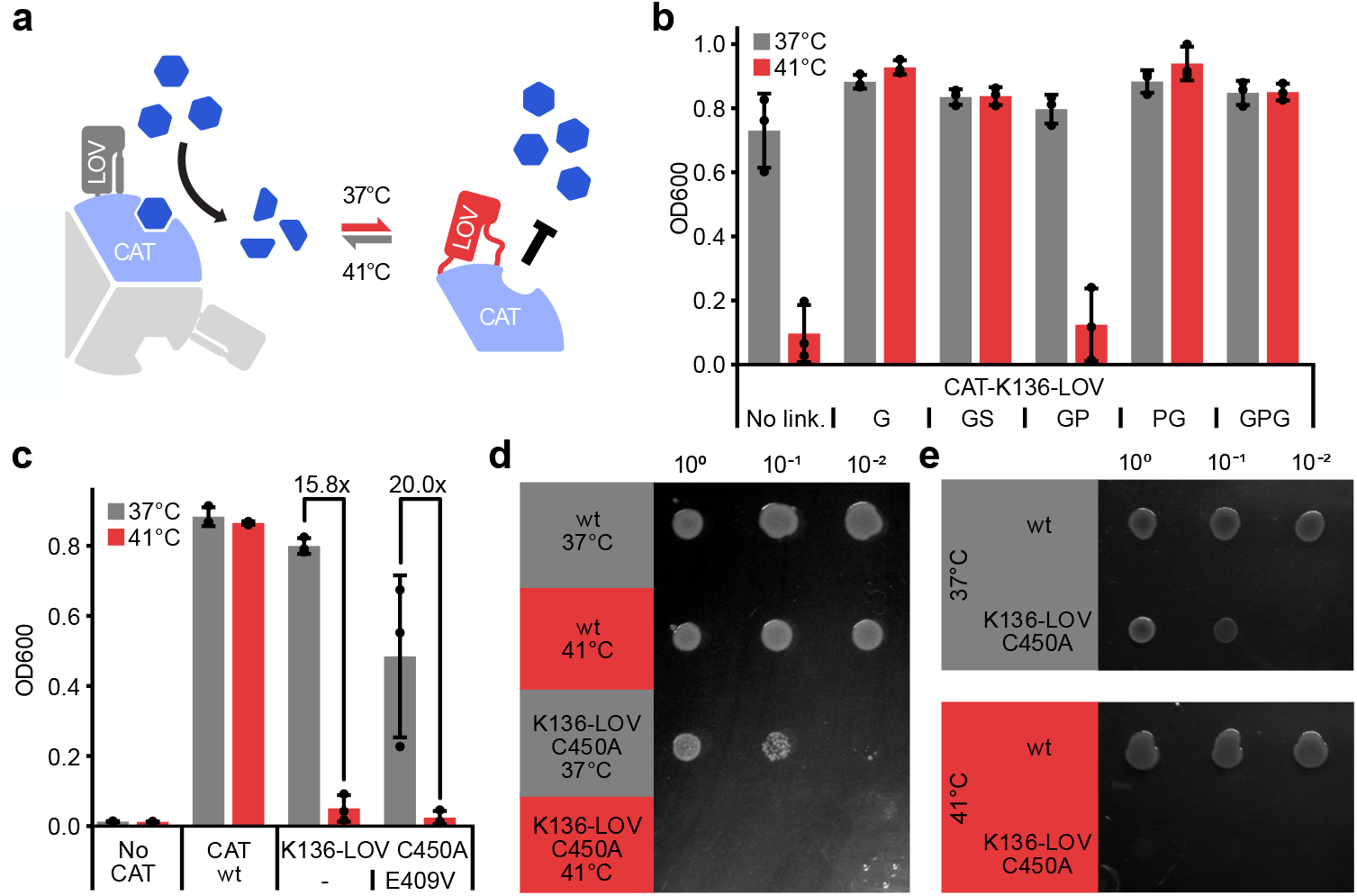
An engineered CAT-LOV hybrid acts as a heat inducible microbial kill switch. **a**, Assay schematics. **b**,**c**, *E. coli* cultures expressing either wild-type CAT, no CAT, the indicated CAT-K136-LOV linker variants (**b**) or CAT-K136-LOV variants with point mutations in the LOV domain (**c**) were grown in the presence of chloramphenicol. Samples were incubated at either 37°C or 41°C for 16 hours, followed by measurement of the OD600. **d**, Serial dilutions of the cultures from **c** (previously incubated at either 37°C or 41°C) were spotted onto LB-agar supplemented with chloramphenicol and incubated at 37°C overnight before the image was taken. **e**, Serial dilutions of pre-cultures expressing either wt CAT or CAT-K136-LOV C450A were spotted onto LB-agar supplemented with chloramphenicol and the plates were incubated overnight at the indicated temperature. **b**,**c**, Data represent the mean +/-SD, n = 3 independent experiments. wt, wild-type.

To apply our method to an additional structurally unrelated protein in *E. coli*, we focused on a *Streptococcus pyogenes* (*Spy*)Cas9 CRISPRa system, in which the transcriptional activator SoxS is recruited to a modified sgRNA scaffold via the MS2 aptamer-binding domain MCP^13,14^ (Supplementary Fig. 8a). We inserted the *As*LOV2-C450A domain into MCP at three different sites and tested these variants at 37°C using RFP expression as a readout. A hybrid carrying the LOV insertion after N27 resulted in strong reporter expression (Supplementary Fig. 8b) and was selected for further characterization. Building on these results, we repeated the CRISPRa experiment at 37°C and 41°C and included the K488N *As*LOV2 mutant, which had shown reduced leakiness in the context of AraC (Supplementary Fig. 8c). As expected, the LOV insertion variants were effectively inactivated at 41°C, while the constitutively active system based on wild-type MCP was only slightly affected by the temperature increase. Consistent with our previous results, the K488N mutant reduced leakiness in the off state (41°C), albeit at the expense of weaker activity at 37°C (Supplementary Fig. 8c). Importantly, our thermogenetically regulated MCP could be easily adapted to the various applications of the MS2 aptamer system such as mRNA tagging^15^ and purification^16^ in a plug-and-play manner.

In addition to allosteric protein control via *As*LOV2 domain insertion, we investigated the conditional caging of signaling peptides in the AsLOV2 J? helix. The light-dependent exposure of functional peptides caged within the C-terminal Jα helix is a commonly used strategy in optogenetics^17–20^. In this case, the peptide is appended to or integrated into the Jα helix, resulting in peptide caging in the *As*LOV2 dark-adapted state due to tight association of Jα with the LOV protein core. In turn, light-induced unfolding of the Jα helix exposes the peptide. To assess whether this concept could be adapted for thermoregulation, we fused RFP to several modified variants of *As*LOV2 in which the C-terminus of the Jα helix was modified to resemble an SsrA degradation tag based on a previously reported design^19^ (Supplementary Fig. 9a). Constitutive expression of these constructs in *E. coli* revealed a strong heat-dependent decrease in RFP levels at 41°C and even more pronounced at 42°C, while RFP expression in controls lacking the LOV-caged SsrA peptide remained unaffected by the temperature condition (Supplementary Fig. 9b).

Collectively, these data demonstrate the general applicability of the *As*LOV2 insertion strategy for temperature-dependent allosteric protein regulation and peptide caging.

### Thermo-control of CRISPR effectors in mammalian cells

Thermogenetic gene editors responding to narrow temperature changes within the human physiological range could fuel various applications in biomedical research and, eventually, precision gene therapy. Thus, we sought to transfer our concept to CRISPR-Cas9 and human cells. First, to create temperature-activated genome editors, we build upon the CASANOVA strategy for Cas9 control with light-switchable anti-CRISPR (Acr) proteins previously developed by us. We inserted *As*LOV2 C450A-(K488N) after position E76 into the Cas9 inhibitor AcrIIA5^21,22^ (Fig. 4a) and co-expressed the resulting AcrIIA5-LOV hybrid with *Staphylococcus aureus* (*Sau*)Cas9 and an sgRNA targeting the endogenous *EMX1* or *GRIN2B* locus in HEK293T cells. We observed an up to 3.4-fold increase in InDel rates at the targeted site after 72 h of incubation at 40°C versus 37°C, as assessed by deep amplicon sequencing. In control samples expressing wild-type AcrIIA5 or no Acr at all, InDel rates were low (<5%) or high (∼60-75%), irrespective of the temperature applied (Fig. 4b-c).

**Figure 4.**
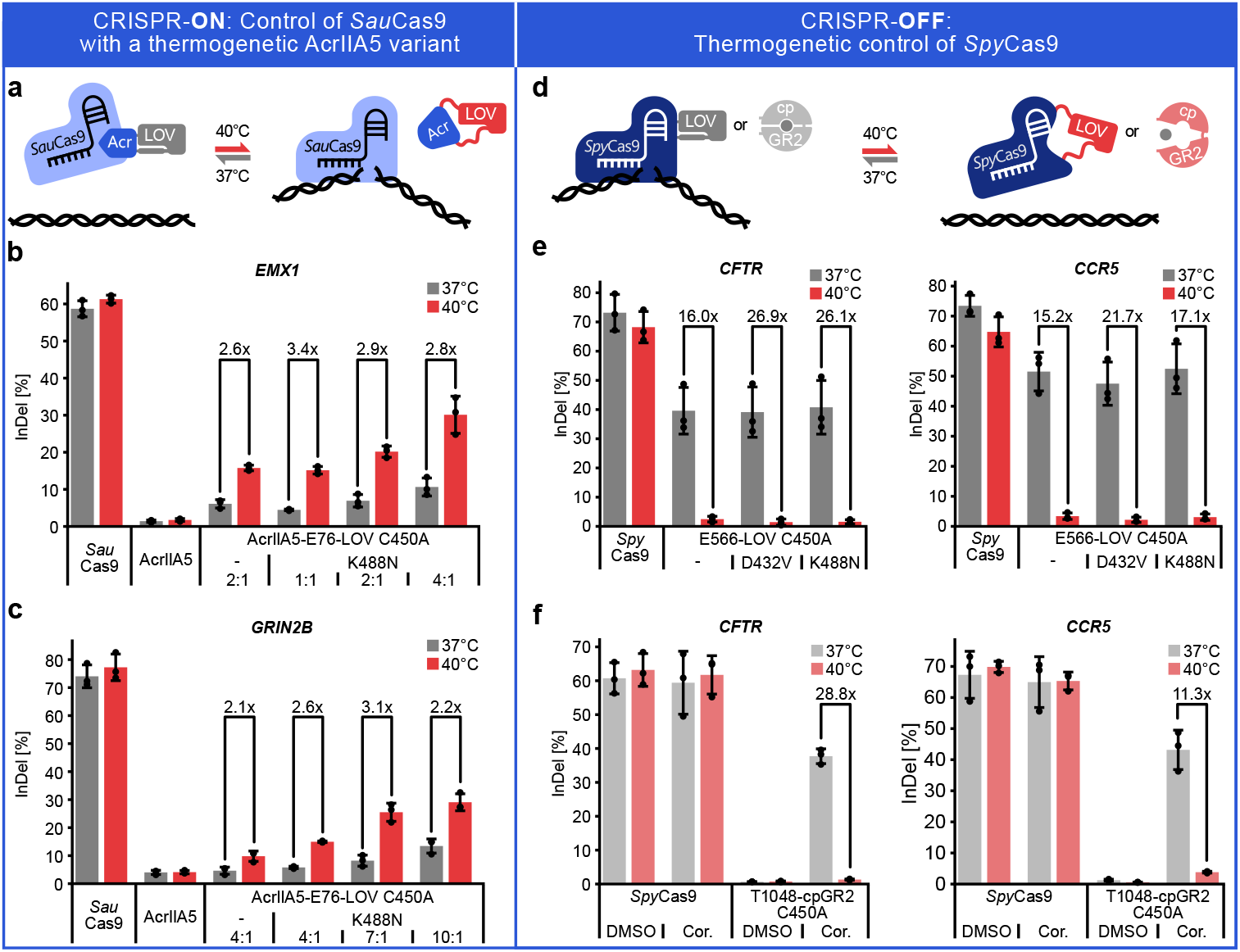
Engineering of potent thermally regulated CRISPR effectors. **a**,**d**, Assay schematics. **b-c**,**e**-**f**, HEK293T cells were transiently transfected with plasmids encoding *Sau*Cas9 (**b**,**c**) or the indicated *Spy*Cas9 variant (**e**,**f**), an sgRNA targeting the indicated endogenous locus. In b-c the respective AcrIIA5(−LOV) variant or an empty stuffer plasmid was co-transfected at the indicated Cas:Acr vector mass ratios. Samples were incubated at 37°C or 40°C and editing efficiency was assessed by deep amplicon sequencing 72 hours post-transfection. **b**-**c**,**e-f**, Data are mean +/-SD, n = 3 independent experiments.

To invert this mechanism of action and demonstrate control of a second Cas9 ortholog, we directly inserted *As*LOV2 variants into *Spy*Cas9 at two allosteric sites (E566 and T1048) that we have recently identified in the context of light-mediated regulation^12^ (Fig. 4d). Both Cas9-LOV hybrids mediated potent, heat-inactivated editing (Supplementary Fig. 10) with the *Spy*Cas9-E566-LOV-C450A hybrid and its *As*LOV2 D432V and K488N point mutants resulting in particularly effective thermo-control across multiple target sites (Fig. 4e and Supplementary Fig. 11a). Remarkably, Cas9 thermoregulation could be plug-and-play applied for CRISPR activation (CRISPRa) as well, simply by inserting LOV mutants at E566 into the dCas9-VPR transcriptional activator (Supplementary Fig. 11b).

Finally, we hypothesized that temperature-sensitivity may not be a specific feature of LOV domains only, but rather a widespread phenomenon of sensory domains. Sensory domains need to be capable of transitioning between different conformational states and may become conformationally unstable when temperature rises above their native working temperature (Supplementary Discussion). To investigate this hypothesis experimentally, we selected a circularly permuted glucocorticoid receptor (cpGR2) domain recently reported by us, which adopts a more compact conformation upon cortisol addition^14^. We inserted cpGR2 into *Spy*Cas9 at positions E566 and T1048 and tested the resulting hybrid effectors for temperature- and ligand- (co-)dependent genome editing in HEK293T cells. In the presence of cortisol, both Cas9-cpGR2 hybrids mediated potent *CCR5* editing at 37°C. In contrast, no editing was observed at 40°C, irrespective of cortisol addition (Supplementary Fig. 12). The T1048 variant showed a particularly strong thermal control of genome editing across target loci with up to 28-fold changes between the temperature conditions, whereas control samples remained unaffected (Fig. 4f). These experiments demonstrate that alike *As*LOV2, cpGR2 in its cortisol-induced state, is inherently temperature-sensitive.

## Discussion

We introduced the modular engineering of allosteric thermogenetic protein switches by receptor domain insertion as a powerful method for tailored protein regulation and demonstrated its applicability to diverse effectors in both bacteria and mammalian cells. The success of this approach directly raises the question of whether the chosen sensory domains are, by accident, particularly thermosensitive or whether this is a widespread property of sensory domains in nature. To date only very few proteins or domains are known to undergo changes in activity and/or conformation in response to small temperature changes close to the human physiological temperature optimum^23^. Of note, another recent thermogenetic study also built on a (photo)receptor domain, i.e. *Bc*LOV4, to control membrane recruitment of proteins^8^. We speculate that the apparent suitability of sensory receptor domains for thermo-regulation may be explained by their specific structural and functional properties. In order to respond to any kind of stimulus, sensory domains must be capable of (reversibly) adopting different conformations. From a conformational energy landscape perspective, this means that the stimulus shifts the conformational equilibrium between at least two very different structural states^24^. The *As*LOV2 domain, for instance, is well-known to co-exist in a dark- and light-adapted state, but light induction strongly shifts the equilibrium towards the light-adapted conformation^25^. Importantly, the thermodynamic conformational equilibrium inherently depends on temperature. In other words: Temperature changes can profoundly reshape the conformational energy landscape. This may explain why conformationally promiscuous receptor domains, especially those undergoing order-disorder transitions, may be particularly well-suited for adaption as thermosensors.

In addition, the domain insertion strategy used here imposes structural strain on both the receptor domain and the effector protein. While usually not favorable for opto- or chemogenetic applications, the resulting destabilization of the hybrid protein is likely beneficial for the engineering of protein thermosensors, where a structural change is intended to occur at the low living temperatures of most biological systems^26^. This insight not only opens exciting possibilities for modularly engineering thermogenetic tools by leveraging existing opto- or chemogenetic switches or protein models^27^, but also underscores the importance of considering temperature effects in experimental design across biological research.

From an application perspective, the thermo-regulation of CRISPR tools is of great biomedical interest, but only few approaches for CRISPR-Cas effector control have been reported. While Cas9 orthologs with different thermal stabilities are known^28^, the effect of temperature changes within the physiological range on CRISPR-Cas activity tends to be negligible^29^. Similarly, the only well-characterized naturally thermosensitive Cas9 inhibitor, AcrIIA2, requires large temperature changes, i.e. between 22°C and 37°C, which is not easily compatible with mammalian cells^30^. The only example of a directly thermo-controlled Cas9 was reported by the group of Andreas Möglich and is a hybrid of *Spy*Cas9 and the LOV domain from *Rhodobacter sphaeroides* (*Rs*LOV)^31^. Of note, this variant, which has only been used in bacteria, still requires a relatively large temperature shift from 29°C to 37°C. Finally, a common workaround represents the thermo-controlled expression of CRISPR-Cas using heat-shock promoters or delivery via photothermally regulated nanoparticles^6,32–34^. These systems, however, require several components, cannot be combined with e.g. cell type-specific promoters, and heat-shock promoters are prone to crosstalk with other cellular processes.

Our system, in contrast, combines several highly advantageous features, including precise response within the small physiological temperature window, direct control of Cas9 rather than indirect transcriptional regulation, tunability of the transition temperature, and the ability to provide both heat-activated and heat-inactivated control of genome editing by integrating *As*LOV2 into either Cas9 or an Acr. In addition, our strategy is modular with regard to the promoter used for expression, e.g. it would be compatible with cell type-specific promoters, and should even be compatible with non-DNA-based delivery strategies (mRNA). In light of these advancements, we strongly believe that our method solves several long-standing challenges regarding thermo-regulation of proteins.

## Methods

### Molecular cloning

The plasmids generated and used in this study are listed in Supplementary Table 1 and the corresponding genbank files are provided in Supplementary Data 1. The amino acid sequences of the relevant proteins are shown in Supplementary Table 2 and the engineered variants and mutants thereof are shown in Supplementary Figure 3. The constructs were generated either by Golden Gate assembly, Gibson assembly or by kinase, ligase and DpnI (KLD) treatment. Linker sequences and type IIS restriction enzyme recognition sites were introduced as 5’-primer overhangs. Oligonucleotides were purchased from Merck, and PCRs were performed using Q5 high-fidelity DNA polymerase (New England Biolabs) under standard conditions. PCR products were analyzed on agarose gels, and the correct bands were excised and purified using the QIAquick gel extraction kit (Qiagen). Assembly reactions were performed using enzymes and buffers purchased from New England Biolabs and Thermo Fisher Scientific. Chemically competent Top10 *E. coli* cells (Thermo Fisher Scientific) were transformed with the assembled constructs and plated on agar supplemented with the appropriate antibiotic. Finally, plasmid DNA was purified from the liquid cultures using the QIAprep Spin Miniprep or Plasmid Plus Midi kit (Qiagen). All constructs were verified by Sanger sequencing using Microsynth Seqlab. The *Renilla* luciferase expression plasmid was purchased from Promega. pX601-AAV-CMV::NLS-SaCas9-NLS-3xHA-bGHpA;U6::BsaI-sgRNA was a gift from Feng Zhang (Addgene plasmid # 61591 ; http://n2t.net/addgene:61591 ; RRID:Addgene_61591) and pEJS654 All-in-One AAV-sgRNA-hNmeCas9 was a gift from Erik Sontheimer (Addgene plasmid #112139; http://n2t.net/addgene:112139; RRID: Addgene_112139). sgRNA1_Tet-inducible Luciferase reporter and Tet-inducible mCherry reporter were a gift from Moritoshi Sato (Addgene plasmid #64161, #64128 ; http://n2t.net/addgene:64161 ; RRID:Addgene_64161)

### Thermal control setup

All experiments in which samples were cultured at up to three different temperatures were performed in standard shaking incubators for bacteria or humidified incubators for mammalian cells. The incubators were operated with identical settings except for temperature, which was adjusted as indicated. Temperature settings were validated periodically using a Xintest HT-9815 thermometer and K-type thermocouples.

### AraC reporter assays

Pre-cultures of *E. coli* Top10 cells transformed with constructs encoding the respective AraC variant and the mRFP reporter were inoculated from glycerol stocks and cultured overnight in 4 mL LB media supplemented with chloramphenicol (25 µg/mL) and kanamycin (50 µg/mL) at 37 °C and 220 rpm. The next day, main cultures were prepared by inoculating LB media supplemented with chloramphenicol (25 µg/mL), kanamycin (50 µg/mL), 400 µM IPTG and 16 mM arabinose (unless otherwise specified) with the pre-cultures. Cultures were grown for 16 hours before mRFP fluorescence and OD600 were measured in a Tecan infinite 200 plate reader. Alternatively, RFP levels of 20,000 cells were measured in a Cytoflex S flow cytometer (Beckman Coulter). The gating strategy is shown in Supplementary Fig. 13a-b. Initial experiments and characterization of the thermo-responsive variants were performed in 15 mL culture tubes using 4 mL media inoculated with 40 µL of a pre-culture. Dose escalation and reversibility assays were performed in 48-well plates in a 500 µL culture volume inoculated with 5 µL of pre-culture.

To measure reversibility of reporter expression, pre-cultures were diluted 1:30 in 500 µL of LB media supplemented with all antibiotics and inducers in 48-well plates and incubated at 37°C or 41°C for 5 hours before they were moved to the other temperature condition, respectively, and incubated for another 19 hours. After 3 hours and 7 hours of incubation, the samples were diluted 1:20 and 1:30 in fresh media. RFP fluorescence and OD600 were measured at the beginning of the experiment and after 3 hours, 7 hours and 24 hours, respectively.

For simultaneous measurements at different temperatures, samples were incubated along a heat gradient in a thermal cycler as previously described by Piraner et al. ^4^. Pre-cultures were diluted to an OD600 of 0.25, inducers (400 µM IPTG and 16 mM arabinose) were added and 25 µL aliquots of the culture were divided into 12 0.2 mL PCR tubes. The tubes were placed in a thermal cycler (Eppendorf) and the gradient function was used to incubate each sample at a different temperature. After 18 hours of incubation, 75 µL of LB medium was added to each sample and 90 µL were transferred to a 96-well microtiter plate. Fluorescence measurements were performed as described above. To infer transition temperatures, the mean reporter values for each AraC-LOV variant were min-max normalized and fitted to the Hill equation using the Neutcurve python package^35^.

To evaluate the spatial control of gene expression, 2.5% LB agar supplemented with chloramphenicol (25 µg/mL), kanamycin (50 µg/mL), 400 µM IPTG and 16 mM arabinose was prepared. Bacteria from pre-cultures were streaked onto the agar, which was then placed in a metal tray, half of which was heated to ∼41°C for 16 hours. Images were captured under blue light.

### Directed evolution experiments

To create an *As*LOV2 mutant library, the LOV domain was amplified by error-prone PCR using the GeneMorph II Random Mutagenesis Kit (Agilent) according to the manufacturer’s protocol for medium mutation rates. The AraC expression plasmid was opened at position S170 by around-the-horn PCR using Q5 Hot Start high-fidelity DNA polymerase (New England Biolabs). The primers carried type IIS restriction enzyme recognition sites as overhangs enabling the efficient assembly of the AraC-S170-LOV fusion sequence through Golden Gate cloning. After Golden Gate assembly, electrocompetent *E. coli* Top10 cells carrying a pBAD-driven mRFP reporter were transformed with the plasmid library using a Gene Pulser Xcell (Biorad) at 1800 V and 200 Ω. The transformed cells were allowed to recover for 1 hour in 1 mL SOC medium without antibiotics while incubated at 37°C and 800 rpm in a thermoshaker (Eppendorf). Next, a 10 mL culture of LB medium supplemented with chloramphenicol (25 µg/mL) and kanamycin (50 µg/mL) was inoculated with the whole 1 mL culture and grown overnight. In parallel, serial dilutions of the transformants were plated on agar and used to determine a transformation efficiency of 5.2 million colony forming units. Sanger sequencing of randomly picked colonies revealed an amino acid substitution rate of ∼1.3 per clone. Considering a theoretical library complexity of 2,679 possible single amino acid mutants and 7.2 million double mutants, all single mutants and many of the possible double mutants were covered. Finally, glycerol stocks of the library were prepared in aliquots and stored at -80°C until further use.

To select the library for thermo-responsive candidates, a pre-culture was grown from the glycerol stocks overnight at 37°C and 220 rpm. The next day, the cultures were diluted 1:1,000,000 and 400 µL were plated onto 20 large LB agar petri dishes (20 cm diameter) supplemented with chloramphenicol (25 µg/mL), kanamycin (50 µg/mL), 400 µM IPTG and 16 mM arabinose. Plates were incubated overnight at 40°C and photographed under blue light. The plates were then incubated at 37°C for an additional 2 hours and imaged again. Next, pictures were analyzed computationally using an modified version of the “colony counter” package (https://github.com/morris-lab/Colony-counter). Colonies that were non-fluorescent after the initial incubation at 40°C, but showed bright fluorescence after the 37°C incubation step were considered promising candidates. These clones were picked, and their performance was tested using the quantitative reporter assay described above. Mutations in the verified lead candidates were determined via Sanger sequencing.

### Antibiotic resistance assays

Pre-cultures of *E. coli* Top10 cells transformed with the respective CAT-encoding construct were inoculated from glycerol stocks and grown overnight in 4 mL LB media supplemented with ampicillin (100 µg/mL) at 37°C and 220 rpm. The next day, main cultures were prepared by adding 5 µL pre-cultures to 48-well plates containing 500 µL of LB media supplemented with 100 µg/mL ampicillin and 25 µg/mL chloramphenicol in technical duplicates. Two plates were prepared, one incubated at 37°C and 220 rpm, the other at 41°C and 220 rpm. After 16 hours of incubation, the optical density (OD) was measured at 600 nm in a Tecan infinite 200 plate reader. The values of an LB-only control were subtracted from each sample. For quantitative assessment of cell survival, serial dilutions of main cultures grown at different temperatures were subsequently spotted onto LB agar supplemented with 100 µg/mL ampicillin and 25 µg/mL chloramphenicol and grown overnight at 37°C before images were taken. Thermally induced killing of bacteria growing on agar was performed in a similar manner. In this case, serial dilutions of a pre-culture were spotted onto two separate replicate LB agar plates supplemented with 100 µg/mL ampicillin and 25 µg/mL chloramphenicol, one incubated overnight at 37°C and the other at 41°C.

To measure growth curves, cultures were prepared identically but incubated in the plate reader at 37°C for 16 hours, while shaking at a 4 mm radius. The OD600 was measured every 15 minutes.

### Inducible protein degradation

Top10 *E. coli* cells were transformed with plasmids encoding different mRFP-LOV fusion reporter proteins. Pre-cultures were inoculated from glycerol stocks and grown overnight at 37°C and 220 rpm in 4 mL LB media supplemented with (50 µg/mL) kanamycin. The next day, three identical main cultures were prepared, by inoculating 4 µL of the respective pre-cultures into 4 mL of LB media supplemented with the same antibiotic and 1 mM arabinose, followed by incubation at 220 rpm and 37°C, 41°C or 42°C, respectively. After 16 hours of incubation, mRFP fluorescence and OD600 were measured in a Tecan infinite 200 plate reader.

### SoxS-mediated CRISPRa in *E. coli*

Top10 *E. coli* cells were co-transformed with plasmids encoding (i) an mRFP reporter and (ii) d*Spy*Cas9, an sgRNA with an MS2 stem loop integrated into its scaffold and a MCP-SoxS fusion protein or variants thereof. Pre-cultures were inoculated from glycerol stocks and grown overnight at 37°C and 220 rpm in LB media supplemented with 100 µg/mL ampicillin and 25 µg/mL chloramphenicol. The next day, two identical main cultures were prepared, by inoculating 4 mL of LB media supplemented with the same antibiotics with 100 µL of the respective pre-cultures, followed by incubation at 220 rpm and 37°C or 41°C, respectively. After 16 hours of incubation, mRFP fluorescence and OD600 were measured in a Tecan infinite 200 plate reader. The values of an LB only control were subtracted from each sample.

### Human cell culture

HEK293T cells were maintained in 1x DMEM without phenol red (Thermo Fisher Scientific) supplemented with 10% (v/v) fetal bovine serum (Thermo Fisher Scientific), 2 mM L-glutamine, 100 U/mL penicillin, and 100 μg/mL streptomycin (all from Thermo Fisher Scientific). Cells were cultured at 37°C with 5% CO2 in a humidified incubator and passaged at a confluency of 70-80%. Authentication of the cell line was performed before use, and routine testing for mycoplasma contamination was conducted. Cell growth at 40°C was visually inspected with a Keyence BZ-9000 microscope on a regular basis (Supplementary Fig. 14).

### Transient transfection

HEK293T cells were seeded at a density of 12,500 cells per well in 100 µL of media into a 96-well plate. After 24 hours, cells were transfected with a total of 150 ng DNA using 0.5 µL Lipofectamine 2000 (Thermo Fisher Scientific) per well according to the manufacturer’s instructions. For genome editing using *Spy*Cas9, cells were co-transfected with (i) 75 ng of a construct expressing either wild-type *Spy*Cas9 or a *Spy*Cas9-LOV or *Spy*Cas9-GR2 hybrid and (ii) 75 ng of the corresponding sgRNA expression vector. For AcrIIA5-dependent genome editing with *Sau*Cas9, cells were co-transfected with 150 ng (i) of an all-in-one construct encoding *Sau*Cas9 and an *EMX1* or *GRIN2B* targeting sgRNA and (ii) AcrIIA5 wild-type or the AcrIIA5-E76-LOV2 mutant at the vector mass ratios indicated in the figures. Cas9 without sgRNA was transfected as a negative control. The sgRNA sequences used in this study are listed in Supplementary Table 4. After transfection, all samples were either incubated for 72 h at 37°C or 40°C with 5% CO2 in a humidified incubator. For the chemical induction, 1 µl of 20 µM Cortisol was added per well after 2 hours, whereby 30 mM of cortisol dissolved in DMSO was pre-diluted 10-fold in DMEM.

### Genome editing of endogenous loci

Editing efficiency at endogenous loci was assessed by deep amplicon sequencing or T7 endonuclease I (T7EI) assay, as indicated. In both cases, samples were processed 72 h post-transfection. Media were aspirated and cells were lysed in DirectPCR Lysis Reagent (PeqLab) supplemented with 200 µg/mL proteinase K (Roche Diagnostics). Lysis was performed in a shaking incubator at 120 rpm and 55°C for at least 6 hours, followed by heat inactivation of proteinase K at 85°C for 45 minutes. Targeted genomic loci were PCR amplified using Q5 High-Fidelity 2X Master Mix (NEB) with the primers listed in Supplementary Table 5. Dual-barcoded versions of these primers were used for NGS. Correct amplicon length and purity were determined by running samples on a 1% 0.5x Tris-acetate-EDTA (TAE) agarose gel. Barcoded samples were pooled in equimolar ratios and Illumina sequencing was performed by GENEWIZ using the Amplicon-EZ service. Indel frequencies were calculated using CRISPresso 2.0 as previously described (https://github.com/pinellolab/CRISPResso2) ^36,37^. For T7EI assays, 5 µL of PCR amplicons were annealed in 20 µL of 1x NEB Buffer 2 by heating the samples to 95°C for 5 minutes followed by gradual cooling to 25°C. The annealed samples were incubated with 0.5 µL T7EI at 37°C for 15 minutes. Gene editing efficiency was quantified by running samples on a 2% 1x Tris-borate-EDTA (TBE) agarose gel and analyzing band intensities using imageJ. The editing efficiency was calculated as 100*(1-SQRT(1 – cleaved fraction)), where the cleaved fraction is the sum of the intensities of the cleaved bands divided by the total band intensities.

### d*Spy*Cas9-VPR mediated CRISPRa in mammalian cells

HEK293T cells were seeded at a density of 75,000 cells per well in 600 µL of media into a 24-well plate. After 24 hours, cells were transfected with a total of 600 ng DNA using 1.8 µL Lipofectamine 3000 (Thermo Fisher Scientific) according to the manufacturer’s instructions. Cells were co-transfected with (i) 240 ng of d*Spy*Cas-VPR wild-type or d*Spy*Cas-LOV-VPR hybrid, (ii) 120 ng of TetO-targeting sgRNA, (iii) 120 ng of a 13xTetO-mCherry-MODC reporter plasmid co-encoding a constitutively expressed EGFP-MODC as transfection control and (iv) 120 ng stuffer plasmid. Cells were incubated at either 37°C or 40°C for 48 hours. For the flow cytometry assay, the cells were washed with 1x phosphate-buffered saline (PBS), trypsinized and resuspended in 1x DMEM without phenol red supplemented with 10% (v/v) fetal bovine serum, 2 mM L-glutamine, 100 U/mL penicillin, and 100 μg/mL streptomycin (all from Thermo Fisher Scientific). The cells were transferred to 2 ml tubes and kept on ice from this stage onward. The samples were washed with 800 µL PBS, resuspended in 250 µL PBS and passed through a 0.45 µM cell strainer. Flow cytometry was carried out on a Cytoflex S (Beckman Coulter). Cells were identified using the forward scatter and side scatter channels as indicated in Supplementary Figure S13c-d. For each sample, 20,000 cells were recorded. The mCherry fluorescence signal was measured with a yellow laser (561 nm excitation, 610 nm emission). The collected data were subsequently processed and analyzed using the Cytoflow software (https://github.com/cytoflow/cytoflow).

## Data Availability

Additional information, including relevant amino acid sequences, targeted genomic loci and PCR primers for InDel quantification are provided in the Supplementary Information. Genbank files of the DNA constructs are provided as Supplementary Data files. Important constructs will be shared on Addgene (Addgene ID: xxxx, xxxx). Additional data will be shared upon reasonable request.

## Acknowledgements

We thank all members of the Niopek laboratory and the Mathony group for helpful discussions.

## Contributions

J.M. conceived the study and directed the work with support from D.N.. K.H., A.K., N.M., N.L., N.H. and S.A. performed the experiments. K.H., A.K., N.M., N.L., N.H., D.N. and J.M. designed and interpreted the experiments. B.W. predicted the domain insertion sites. N.S. established the CRISPRa assay. A.B. and F.V. performed the initial experiments. J.M. and D.N. secured funding. J.M. wrote the manuscript with support from D.N.. All authors approved the final manuscript.

