## Supplementary Information for "Modular Engineering of Thermo-Responsive Allosteric Proteins"

**Content:**

Supplementary Figure 1-14

Supplementary Table 1-5

Supplementary References

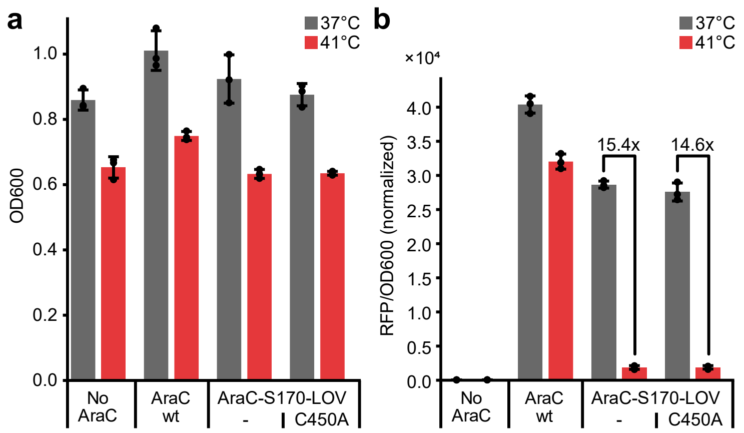


**Supplementary Figure 1 | Insertion of *As*LOV2 into AraC enables thermo-switchable gene expression in *E. coli*. a, b,** *E. coli* containing a pBad-mRFP reporter and expressing the indicated AraC variant or a dummy control protein of similar size were incubated at 37°C or 41°C for 16 h. Culture density (OD600) (**a**) and RFP expression (normalized to OD) (**b**) were assessed in a plate reader. Data points indicate n=3 independent experiments and bars represent the mean. Error bars indicate the standard deviation (SD). Fold changes are indicated. wt, wild-type.

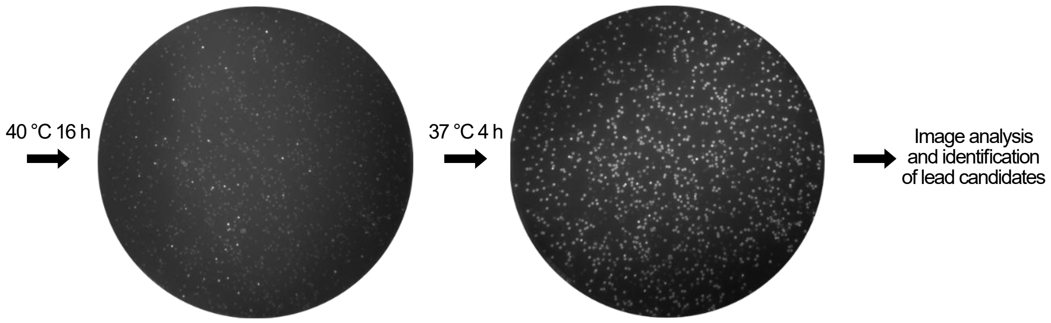


**Supplementary Figure 2 | Selection of improved, thermo-switchable AraC-LOV2 hybrid mutants via screening on agar plates.** Workflow of the AraC-S170-LOV library screening and corresponding agar plate images. Images of the same agar plate were acquired under blue light illumination, first after an initial overnight incubation at 40°C and then after an additional 4 hours at 37°C. Colonies showing no/low fluorescence at 40°C and high fluorescence at 37°C indicate variants with potent photoswitching and were therefore selected for downstream analysis.

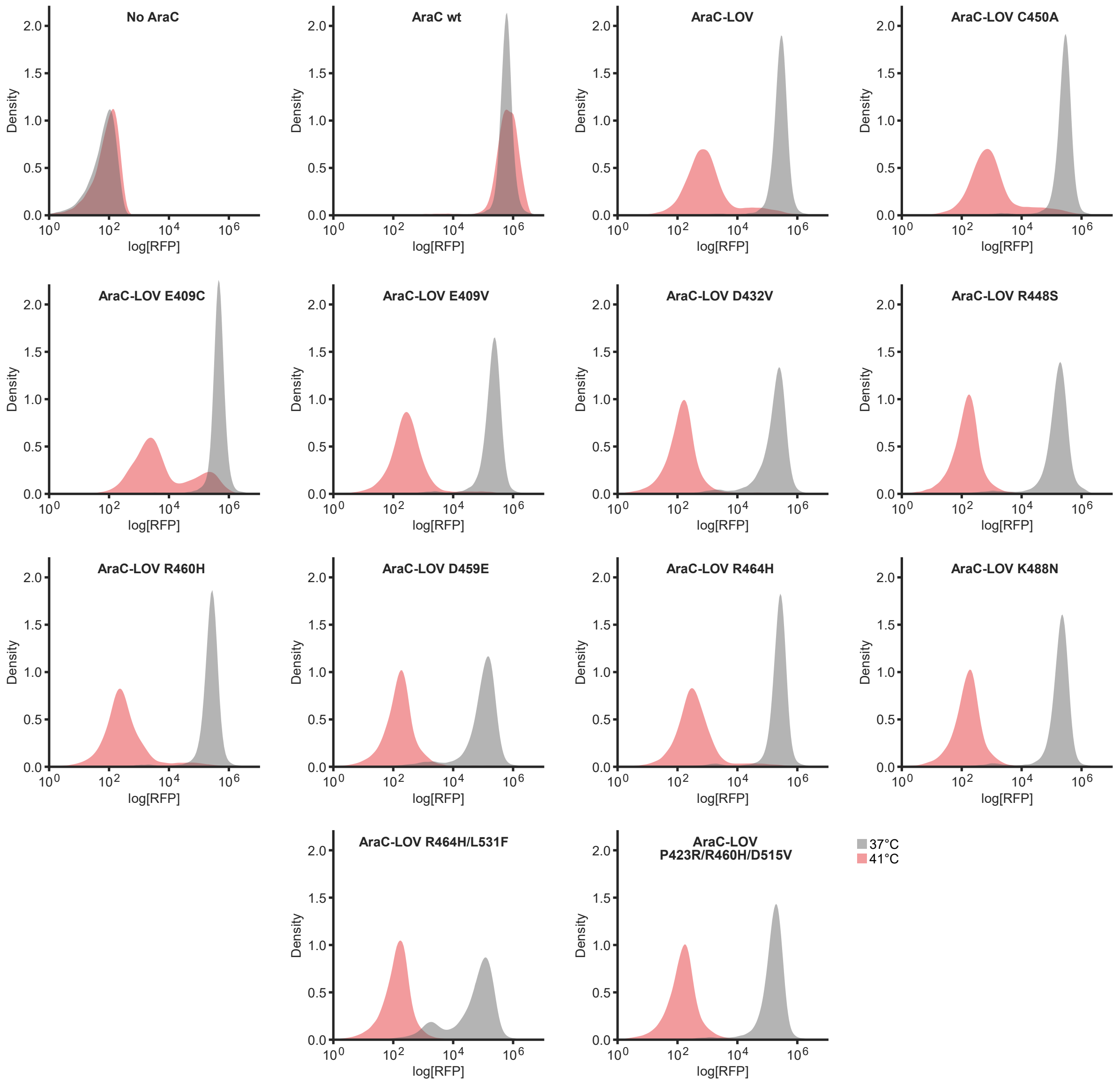


**Supplementary Figure 3 | Directed evolution yields several potent thermosensitive LOV variants.** *E. coli* carrying an AraC reporter and expressing AraC, AraC-LOV, the indicated AraC-LOV-C450A variant or a dummy protein as control were grown at 37°C or 41°C for 16 h, followed by flow cytometry analysis. Histograms show the distribution of fluorescence within the respective sample population for n=3 pooled independent replicates. Wt, wild-type.

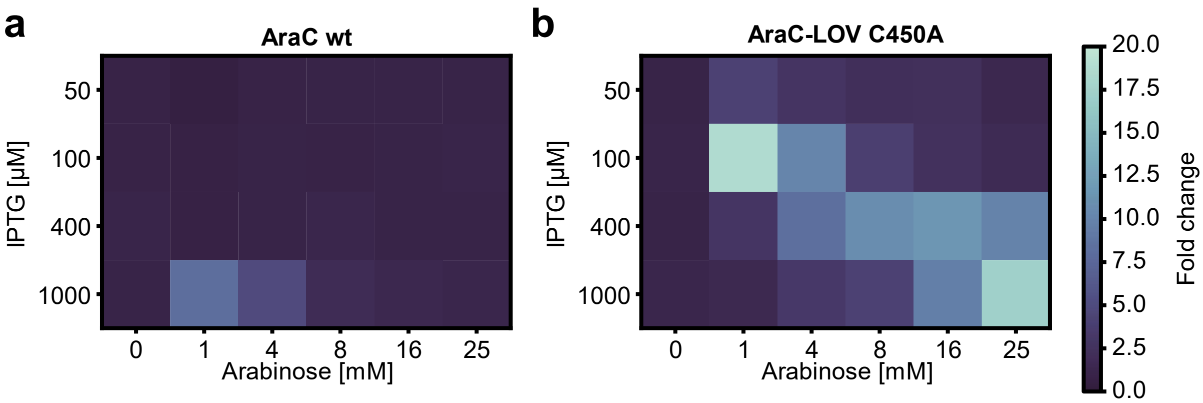


**Supplementary Figure 4 | The thermal response of AraC-LOV can be tuned by varying inducer concentrations. a**, **b**, Fold changes between samples incubated at 37°C and the 41°C condition are shown for *E. coli* expressing either AraC wt (**a**) or AraC-LOV C450A (**b**). Data corresponds to the dose escalation screen in Fig. 2a. wt, wild-type.

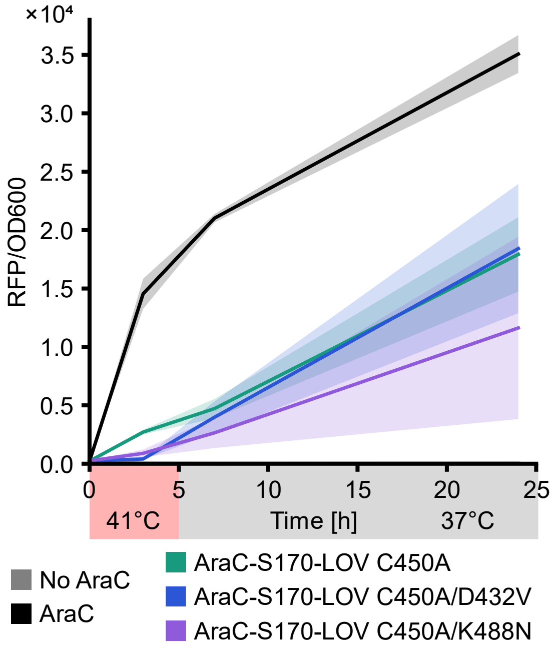


**Supplementary Figure 5 | Temporal control of gene expression.** *E. coli* containing a pBad-mRFP reporter and expressing the indicated AraC variant or a dummy control protein were incubated at 41°C for 5 hours followed by another 19 hours of incubation at 37°C. Culture density (OD600) and RFP expression (normalized to OD) were periodically assessed in a plate reader. Data corresponds to the mean of n=3 independent experiments. The SD is indicated.

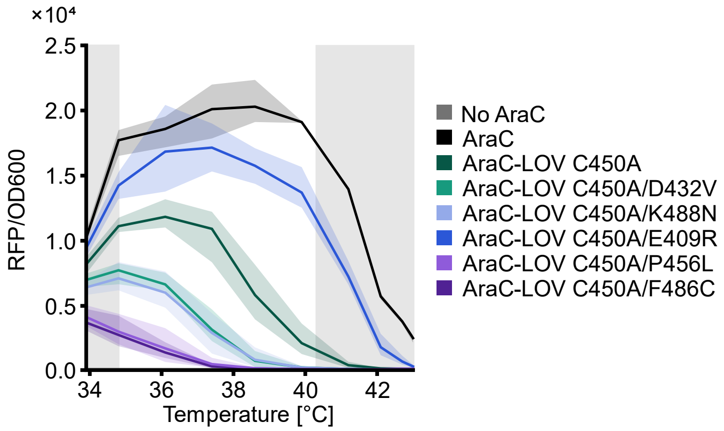


**Supplementary Figure 6 | Point mutations tune the transition temperature and amplitude of thermogenetic AraC.** *E. coli* encoding the pBAD-mRFP reporter and the indicated AraC variant or a control without AraC were incubated at different temperatures between 34°C and 43°C followed by measurement of RFP fluorescence and OD600 in a plate reader. Lines represent the mean of n=3 independent biological replicates. Shaded areas indicate the SD. Regions marked in gray indicate reduced accuracy of the assay due to significant decrease in AraC wild-type reporter levels at these *E. coli* growth conditions.

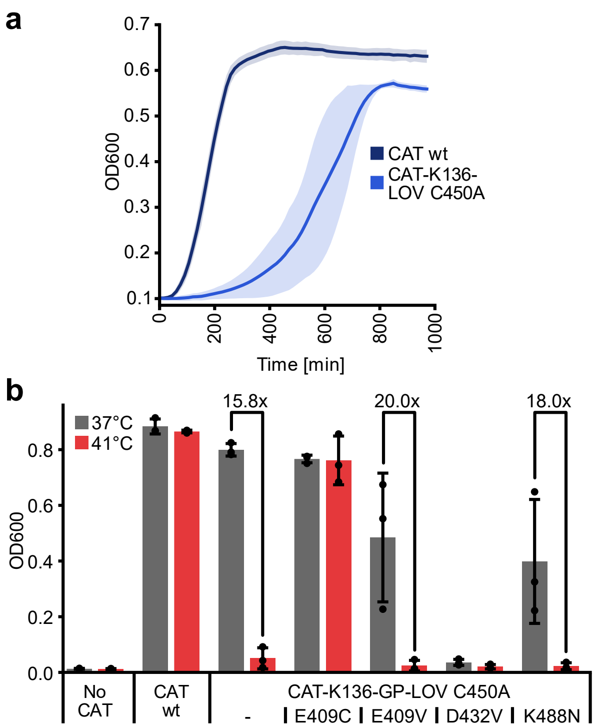


**Supplementary Figure 7 | Characterization of thermosensitive CAT variants. a,** Time course showing the growth of *E. coli* cultures expressing the indicated CAT variants at 37°C in the presence of chloramphenicol. OD600 was measured every 15 minutes for 16 hours using a plate reader. Lines indicate the mean of n=3 independent replicates and shaded areas show the standard deviation. **b**, *E. coli* cultures expressing either wild-type CAT, no CAT, the indicated CAT-K136-GP-LOV mutants were grown in the presence of chloramphenicol. Replicate samples were incubated at either 37°C or 41°C for 16 hours, followed by measurement of the OD600. The graph is an expansion of the dataset shown in Fig. 3c. Data points indicate n=3 independent experiments and bars represent the mean. Error bars indicate the SD. Fold changes are indicated. wt, wild-type.

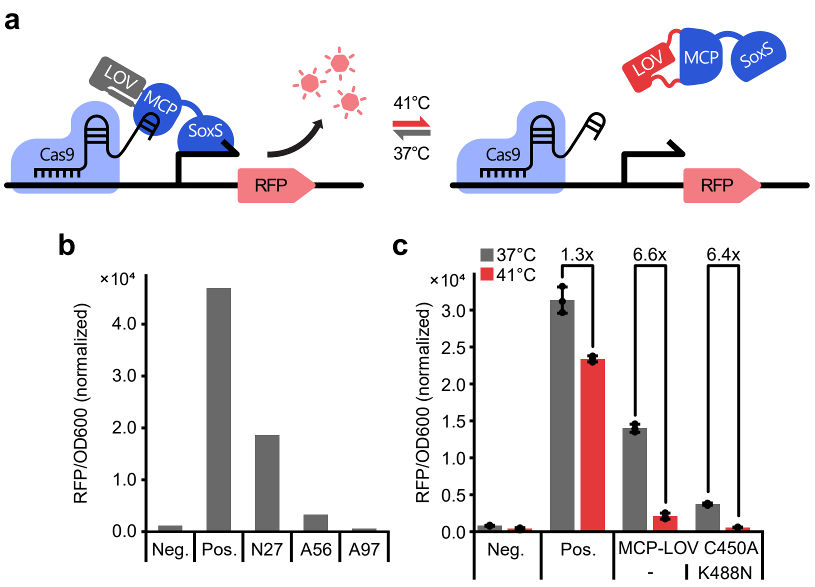


**Supplementary Figure 8 | Screening of LOV insertion sites in MCP. a**, Schematics of MCP-SoxS mediated CRISPRa**. b**, *E. coli* were transformed with plasmids encoding a CRISPRa circuit, i.e. (i) dCas9, (ii) a promoter-targeting or non-targeting sgRNA harboring an MS2 stem loop in its scaffold sequence, (iii) the MCP-SoxS transactivator with an *As*LOV2-C450A insertion after the indicated MCP residue, and (iv) an mRFP reporter. Cultures were incubated at 37°C followed by measurement of mRFP fluorescence and OD600. Bars represent a single experiment. **c**, *E. coli* encoding the same circuit as in b, including the MCP-N27-LOV insertion variant or a K488N point mutant thereof were incubated at 37°C or 41°C for 16 h followed by measurement of mRFP fluorescence and the OD600. Data points indicate n=3 independent replicates and bars represent the mean. Error bars indicate the SD. Fold changes are indicated. Neg., non-targeting negative control; Pos., positive control expressing an mRFP-targeting sgRNA and MCP-SoxS without LOV insertion.

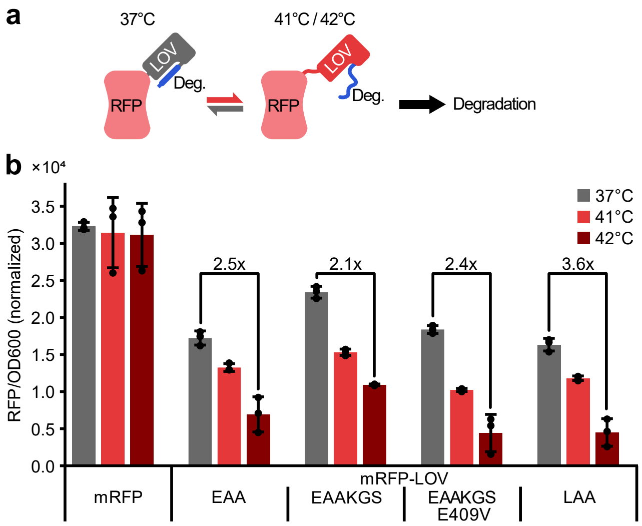


**Supplementary Figure 9 | Temperature-responsive caging of an SsrA degron peptide in the LOV2 Jα helix. a,** Schematic of thermo-switchable RFP degradation. **b**, *E. coli* cultures expressing either mRFP or the indicated mRFP-LOV-degron fusions were grown at the indicated temperatures followed by measurement of mRFP fluorescence and OD600. Letters indicate the C-terminal degron motif starting at amino acid 541 of the *As*LOV2 domain. Data points indicate n=3 independent experiments and bars represent the mean. Error bars indicate the SD. Fold changes are indicated.

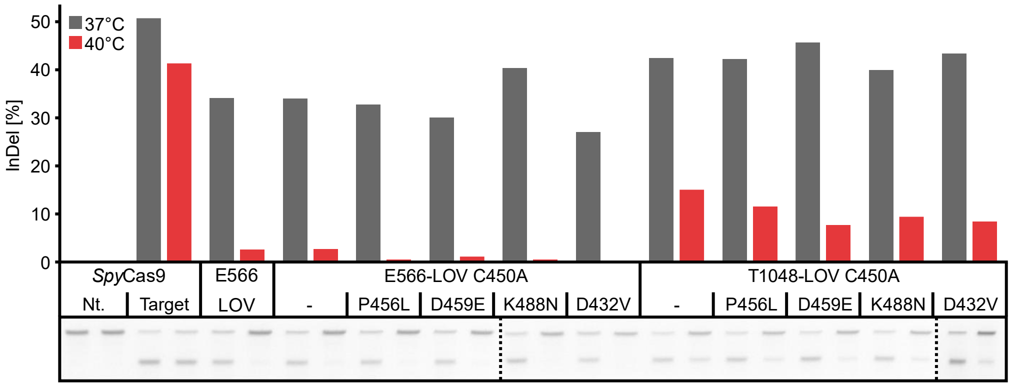


**Supplementary Figure 10 | Screening for thermosensitive *Spy*Cas9 variants.** HEK293T cells were transfected with plasmids encoding (i) an sgRNA targeting the endogenous *CCR5* locus or a non-targeting control and (ii) the indicated Cas9-LOV fusion variant or a wild-type Cas9 control and incubated at 37°C or 40°C. 72 hours post-transfection, editing efficiencies were assessed in a T7EI assay. Bars indicate InDel frequencies from a single experiment, calculated on the basis of the corresponding gel images (bottom). Nt., non-targeting.

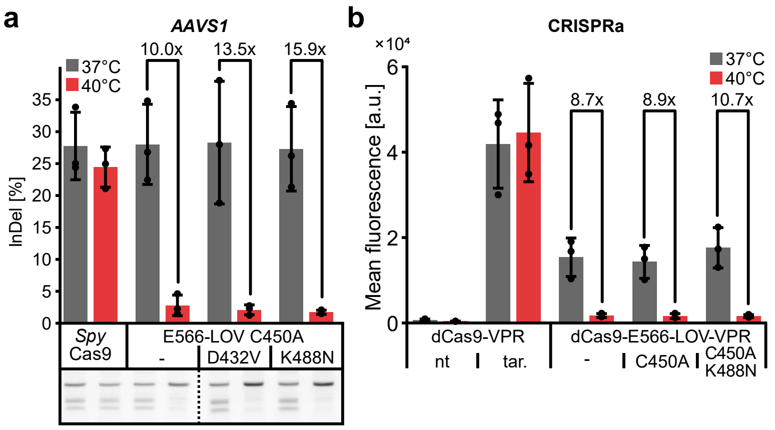


**Supplementary Figure 11 | Targeting AAVS1 with thermo-regulated CRISPR effectors. a,** HEK293T cells were transiently transfected with plasmids encoding (i) wild-type *Spy*Cas9, the indicated *Spy*Cas9 variants with an *As*LOV2 insertion after E566 and (ii) sgRNA targeting the endogenous *AAVS1* locus. Samples were incubated at 37°C or 40°C and editing efficiency was assessed 72 hours post-transfection via T7EI assay. **b**, HEK293T cells were transiently transfected with plasmids encoding (i) d*Spy*Cas9-VPR or the corresponding *As*LOV2 insertion variants, (ii) an mCherry reporter driven from a minimal promoter preceded by 13x TetO repeats, and (iii) a TetO-targeting sgRNA. Samples were incubated at 37°C or 40°C for 48 h before mCherry fluorescence was assessed by flow cytometry. **a**,**b**, Data points of n=3 independent replicates are shown and bars indicate the mean. Error bars indicate the SD. Fold changes are indicated. nt, non-targeting; tar., targeting; a.u., arbitrary units.

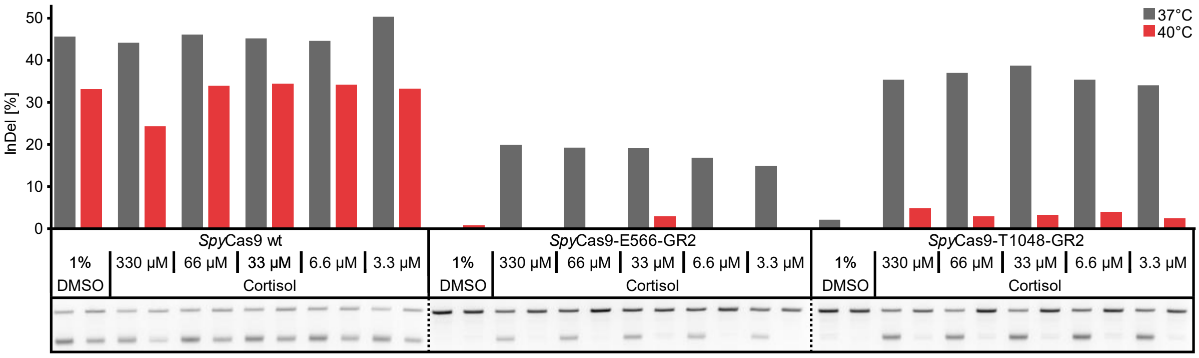


**Supplementary Figure 12 | Cortisol dose screen for *Spy*Cas9-GR2 hybrids.** HEK293T cells were transfected with plasmids encoding (i) an sgRNA targeting the endogenous *CCR5* locus or a non-targeting control and (ii) wild-type Cas9 or a Cas9 variant carrying a GR2 domain insertion after the indicated Cas9 residue. Samples were incubated at 37°C or 40°C and cortisol or DMSO were added 2 hours post-transfection. InDel frequencies were assessed via T7EI assay after 72 hours. Bars indicate InDel frequencies from a single experiment, calculated on the basis of the corresponding gel images (bottom), are shown below. wt, wild-type.

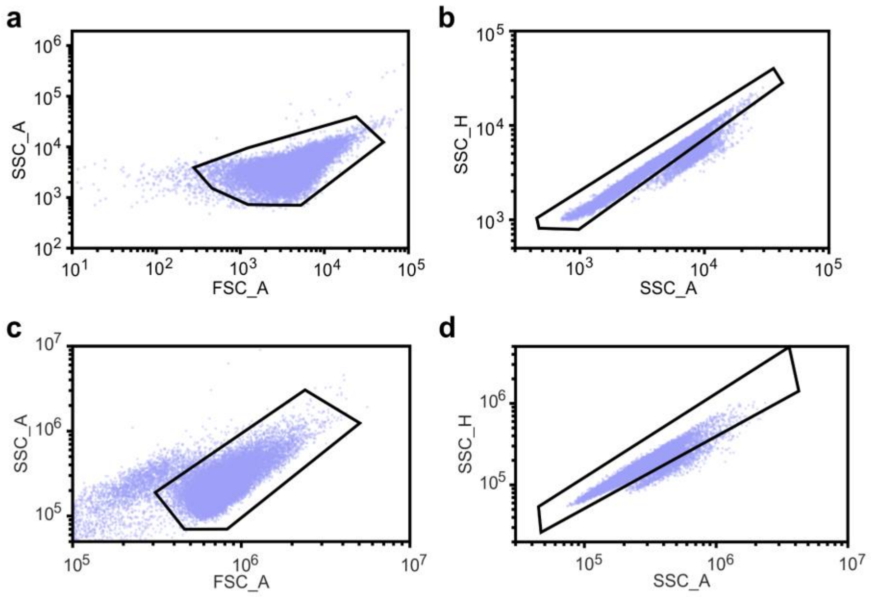


**Supplementary Figure 13 | Gating strategy. a-d**, *E. coli* (**a-b**) or HEK293T cells (**c-d**) were gated based on a forward vs. side scatter area plots (**a**, **c**). Subsequently, single cells were gated within the SSC height and SSC area channels (**b**, **d**).

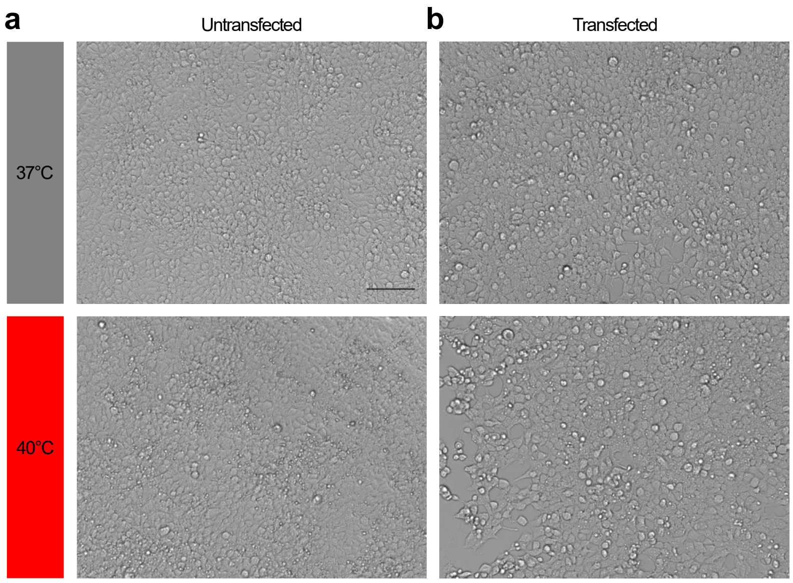


**Supplementary Figure 14 | Cell growth at different temperatures. a,b**, Untransfected (**a**) or transiently transfected (**b**) HEK293T cells were incubated for 72 hours at the indicated temperatures before microscopy images were acquired. Scale bar indicates 100 µM.

**Supplementary Table 1 | List of plasmids used in this study.** Insert domains are flanked by SG linkers on both sides, unless otherwise indicated. CMV, cytomegalovirus; eGFP, enhanced green fluorescent protein; pConst., constitutive promoter.

| # | Name | Description, sequential order | Source |
|---|---|---|---|
| **1** | RFP reporter for AraC | BAD promoter, mRFP1, LVA degradation tag | [1] |
| **2** | AraC | TRC promoter, AraC | [1] |
| **3** | TVMV | TRC promoter, TVMV (negative control) | [1] |
| **4** | BLA | β-lactamase expression cassette | [2] |
| **5** | BLA_CAT | β-lactamase expression cassette, chloramphenicol acetyltransferase expression cassette | [2] |
| **6** | Inducible RFP reporter | Arac, pBAD promoter, mRFP1 | [1] |
| **7** | Inducible RFP-LOVdeg | Arac, pBAD promoter, mRFP1 fused to LOVdegron variant | This study |
| **8** | CRISPRa-RFP-reporter | dCas9-SoxS inducible mRFP | [3] |
| **9** | dspyCas9_MS2_SoxS_non-targeting | pConst, dSpyCas9; J23107 promoter, MCP fused to SoxS via 5xGS linker; J23119 promoter, non-targeting sgRNA with a MS2 stem loop incorporated into its scaffold | [3] |
| **10** | dspyCas9_MS2_SoxS_targeting | pConst, dSpyCas9; J23107 promoter, MCP fused to SoxS via 5xGS linker; J23119 promoter, construct #8-targeting sgRNA with a MS2 stem loop incorporated into its scaffold | [3] |
| **11** | DNA stuffer plasmid | pBluescript sk- | Invitrogen |
| **12** | AcrIIA5 | CMV promoter, wild-type AcrIIA5 from Streptococcus thermophilus, bGHpA | [4] |
| **13** | SauCas9_sgRNA-scaffold | CMV promoter, NLS, SauCas9, NLS, 3xHA, bGHpA; U6 promoter, sgRNA scaffold | [5] |
| **14** | SauCas9_EMX1-sgRNA | CMV promoter, NLS, SauCas9, NLS, 3xHA, bGHpA; U6 promoter, EMX1-targeting sgRNA | [6] |
| **15** | SauCas9_GRIN2B-sgRNA | CMV promoter, NLS, SauCas9, NLS, 3xHA, bGHpA; U6 promoter, GRIN2B-targeting sgRNA | [6] |
| **16** | SpyCas9 | CMV promoter, 3xFlag, NLS, SpCas9, NLS | [7] |
| **17** | CFTR-sgRNA (SpyCas9) | U6 promoter, CFTR-targeting sgRNA, RSV GFP | [7] |
| **18** | CCR5-sgRNA (SpyCas9) | U6 promoter, CCR51-targeting sgRNA, RSV GFP | [8] |
| **19** | AAVS1-sgRNA (SpyCas9) | U6 promoter, AAVS1-targeting sgRNA, RSV GFP | This work |
| **20** | dSpyCas9-VPR | Ef1a promoter, NLS, dSpCas9, NLS, VPR | [9] |
| **21** | mCherry-CRISPRa reporter | TetO repeats, mCherry-MODC, CMV promoter, EGFP-MODC | This work |
| **22** | TetO-sgRNA (SpyCas9) | U6 promoter, TetO-targeting sgRNA | [10] |

**Supplementary Table 2 | Amino acid sequences of the domains and proteins used in this study.** Blue: linker sequences; orange: affinity tag; green: nuclear localization sequence.

| Protein / Domain | Amino acid sequence |
|---|---|
| ***As*LOV2** | LATTLERIEKNFVITDPRLPDNPIIFASDSFLQLTEYSREEILGRNCRFLQGPETDRATVRKIRD<br>AIDNQTEVTVQLINYTKSGKKFWNLFHLQPMRDQKGDVQYFIGVQLDGTEHVRDAAEREGV<br>MLIKKTAENIDEAAK |
| **AraC** | MSAEAQNDPLLPGYSFNAHLVAGLTPIEANGYLDFFIDRPLGMKGYILNLTIRGQGVVKNQGR<br>EFVCRPGDILLFPPGEIHHYGRHPEAREWYHQWVYFRPRAYWHEWLNWPSIFANTGFFRP<br>DEAHQPHFSDLFGQIINAGQGEGRYSELLAINLLEQLLLRRMEAINESLHPPMDNRVREACQ<br>YISDHLADSNFDIASVAQHVCLSPSRLSHLFRQQLGISVLSWREDQRISQAKLLLSTTRMPIAT<br>VGRNVGFDDQLYFSRVFKKCTGASPSEFRAGCEEKVNDVAVKLSGHHHHHH |
| **Chloramphenicol acetyltransferase** | MEKKITGYTTVDISQWHRKEHFEAFQSVAQCTYNQTVQLDITAFLKTVKKNKHKFYPAFIHILA<br>RLMNAHPEFRMAMKDGELVIWDSVHPCYTVFHEQTETFSSLLSEYHDDFRQFLHIYSQDVA<br>CYGENLAYFPKGFIENMFFVSANPWVSFTSFDLNVANMDNFFAPVFTMGKYYTQGDKVLMP<br>LAIQVHHAVCDGFHVGRMLNELQQYCDEWQGGA |
| **RFP-LOVdeg** | MASSEDVIKEFMRFKVRMEGSVNGHEFEIEGEGEGRPYEGTQTAKLKVTKGGPLPFAWDIL<br>SPQFQYGSKAYVKHPADIPDYLKLSFPEGFKWERVMNFEDGGVVTVTQDSSLQDGEFIYKV<br>KLRGTNFPSDGPVMQKKTMGWEASTERMYPEDGALKGEIKMRLKLKDGGHYDAEVKTTYM<br>AKKPVQLPGAYKTDIKLDITSHNEDYTIVEQYERAEGRHSTGASGLATTLERIEKNFVITDPRL<br>PDNPIIFASDSFLQLTEYSREEILGRNARFLQGPETDRATVRKIRDAIDNQTEVTVQLINYTKS<br>GKKFWNLFHLQPMRDQKGDVQYFIGVQLDGTEHVRDAAEREGVMLIKKTAENIDEAA |
| **MCP-SoxS** | MGPASNFTQFVLVDNGGTGDVTVAPSNFANGIAEWISSNSRSQAYKVTCSVRQSSAQNRKY<br>TIKVEVPKGAWRSYLNMELTIPIFATNSDCELIVKAMQGLLKDGNPIPSAIAANSGIYGGGGSM<br>SHQKIIQDLIAWIDEHIDQPLNIDVVAKKSGYSKWYLQRMFRTVTHQTLGDYIRQRRLLLAAVE<br>LRTTERPIFDIAMDLGYVSQQTFSRVFARQFDRTPADYRHRL |
| ***Spy*Cas9** | MDYKDHDGDYKDHDIDYKDDDDKMAPKKKRKVGIHGVPAADKKYSIGLDIGTNSVGWAVITD<br>EYKVPSKKFKVLGNTDRHSIKKNLIGALLFDSGETAEATRLKRTARRRYTRRKNRICYLQEIFS<br>NEMAKVDDSFFHRLEESFLVEEDKKHERHPIFGNIVDEVAYHEKYPTIYHLRKKLVDSTDKAD<br>LRLIYLALAHMIKFRGHFLIEGDLNPDNSDVDKLFIQLVQTYNQLFEENPINASGVDAKAILSAR<br>LSKSRRLENLIAQLPGEKKNGLFGNLIALSLGLTPNFKSNFDLAEDAKLQLSKDTYDDDLDNLL<br>AQIGDQYADLFLAAKNLSDAILLSDILRVNTEITKAPLSASMIKRYDEHHQDLTLLKALVRQQLP<br>EKYKEIFFDQSKNGYAGYIDGGASQEEFYKFIKPILEKMDGTEELLVKLNREDLLRKQRTFDN<br>GSIPHQIHLGELHAILRRQEDFYPFLKDNREKIEKILTFRIPYYVGPLARGNSRFAWMTRKSEE<br>TITPWNFEEVVDKGASAQSFIERMTNFDKNLPNEKVLPKHSLLYEYFTVYNELTKVKYVTEG<br>MRKPAFLSGEQKKAIVDLLFKTNRKVTVKQLKEDYFKKIECFDSVEISGVEDRFNASLGTYHD<br>LLKIIKDKDFLDNEENEDILEDIVLTLTLFEDREMIEERLKTYAHLFDDKVMKQLKRRRYTGWG<br>RLSRKLINGIRDKQSGKTILDFLKSDGFANRNFMQLIHDDSLTFKEDIQKAQVSGQGDSLHEHI<br>ANLAGSPAIKKGILQTVKVVDELVKVMGRHKPENIVIEMARENQTTQKGQKNSRERMKRIEE<br>GIKELGSQILKEHPVENTQLQNEKLYLYYLQNGRDMYVDQELDINRLSDYDVDHIVPQSFLKD<br>DSIDNKVLTRSDKNRGKSDNVPSEEVVKKMKNYWRQLLNAKLITQRKFDNLTKAERGGLSEL<br>DKAGFIKRQLVETRQITKHVAQILDSRMNTKYDENDKLIREVKVITLKSKLVSDFRKDFQFYKV<br>REINNYHHAHDAYLNAVVGTALIKKYPKLESEFVYGDYKVYDVRKMIAKSEQEIGKATAKYFF<br>YSNIMNFFKTEITLANGEIRKRPLIETNGETGEIVWDKGRDFATVRKVLSMPQVNIVKKTEVQT<br>GGFSKESILPKRNSDKLIARKKDWDPKKYGGFDSPTVAYSVLVVAKVEKGKSKKLKSVKELL<br>GITIMERSSFEKNPIDFLEAKGYKEVKKDLIIKLPKYSLFELENGRKRMLASAGELQKGNELAL<br>PSKYVNFLYLASHYEKLKGSPEDNEQKQLFVEQHKHYLDEIIEQISEFSKRVILADANLDKVLS<br>AYNKHRDKPIREQAENIIHLFTLTNLGAPAAFKYFDTTIDRKRYTSTKEVLDATLIHQSITGLYE<br>TRIDLSQLGGDKRPAATKKAGQAKKKKEF |
| ***Sau*Cas9** | MAPKKKRKVGIHGVPAAKRNYILGLDIGITSVGYGIIDYETRDVIDAGVRLFKEANVENNEGRR<br>SKRGARRLKRRRRHRIQRVKKLLFDYNLLTDHSELSGINPYEARVKGLSQKLSEEEFSAALLH<br>LAKRRGVHNVNEVEEDTGNELSTKEQISRNSKALEEKYVAELQLERLKKDGEVRGSINRFKT<br>SDYVKEAKQLLKVQKAYHQLDQSFIDTYIDLLETRRTYYEGPGEGSPFGWKDIKEWYEMLM<br>GHCTYFPEELRSVKYAYNADLYNALNDLNNLVITRDENEKLEYYEKFQIIENVFKQKKKPTLK<br>QIAKEILVNEEDIKGYRVTSTGKPEFTNLKVYHDIKDITARKEIIENAELLDQIAKILTIYQSSEDIQ |

| | |
|---|---|
| | EELTNLNSELTQEEIEQISNLKGYTGTHNLSLKAINLILDELWHTNDNQIAIFNRLKLVPKKVDL<br>SQQKEIPTTLVDDFILSPVVKRSFIQSIKVINAIIKKYGLPNDIIIELAREKNSKDAQKMINEMQKR<br>NRQTNERIEEIIRTTGKENAKYLIEKIKLHDMQEGKCLYSLEAIPLEDLLNNPFNYEVDHIIPRSV<br>SFDNSFNNKVLVKQEENSKKGNRTPFQYLSSSDSKISYETFKKHILNLAKGKGRISKTKKEYL<br>LEERDINRFSVQKDFINRNLVDTRYATRGLMNLLRSYFRVNNLDVKVKSINGGFTSFLRRKW<br>KFKKERNKGYKHHAEDALIIANADFIFKEWKKLDKAKKVMENQMFEEKQAESMPEIETEQEY<br>KEIFITPHQIKHIKDFKDYKYSHRVDKKPNRELINDTLYSTRKDDKGNTLIVNNLNGLYDKDND<br>KLKKLINKSPEKLLMYHHDPQTYQKLKLIMEQYGDEKNPLYKYYEETGNYLTKYSKKDNGPVI<br>KKIKYYGNKLNAHLDITDDYPNSRNKVVKLSLKPYRFDVYLDNGVYKFVTVKNLDVIKKENYY<br>EVNSKCYEEAKKLKKISNQAEFIASFYNNDLIKINGELYRVIGVNNDLLNRIEVNMIDITYREYLE<br>NMNDKRPPRIIKTIASKTQSIKKYSTDILGNLYEVKSKKHPQIIKKGKRPAATKKAGQAKKKKG<br>SYPYDVPDYAYPYDVPDYAYPYDVPDYA |
| **cpGR2** | SGNSSQNWQRFYQLTKLLDSMHEMVGGLLQFCFYTFVNKSLSVEFPEMLAEIISNQLPKFNA<br>GSVKPLLFHQKGGGSGGSGGSGGSGGSGLISLLEVIEPEVLYSGYDSTLPDTSTRLMSTLNR<br>LGGRQVVSAVKWAKALPGFRNLHLDDQMTLLQYSWMSLMAFSLGWRSYKQSNGNMLCFA<br>PDLVINEERMQLPYMYDQCQQMLKISSEFVRLQVSYDEYLCMKVLLLLSTVPKDGLKSQAVF<br>DEIRMTYIKELGKAIVKREGGS |
| **AcrIIA5** | MAYGKSRYNSYRKRNFSISDNQRREYAKKMKELEQAFENLDGWYLSSMKDSAYKDFGKYEI<br>RLSNHSADNRYHDLENGRLIVNVKASKLNFVDIIENKLGKIIEKIDTLDLDKYRFINATKLERDIK<br>CYYKGYKTKKDVI |

**Supplementary Table 3 | Hybrid proteins used in this study.** The insertion site corresponds to the residue in the effector protein preceding the insert domain. The sequences of the two linkers flanking the insert domain are shown.

| # | Name | Protein | Insertion site | Linker | Mutations |
|---|---|---|---|---|---|
| 1 | AraC-S170-LOV | AraC | S170 | SG-GS | - |
| 2 | | | | | C450A |
| 3 | | | | | C450A/E409C |
| 4 | | | | | C450A/E409R |
| 5 | | | | | C450A/E409V |
| 6 | | | | | C450A/D432V |
| 7 | | | | | C450A/R448S |
| 8 | | | | | C450A/D456L |
| 9 | | | | | C450A/D459E |
| 10 | | | | | C450A/R460H |
| 11 | | | | | C450A/R464H |
| 12 | | | | | C450A/S486C |
| 13 | | | | | C450A/K488N |
| 14 | | | | | C450A/R464H/L531F |
| 15 | | | | | C450A/P423R/R460H/D515V |
| 16 | CAT-K136-LOV | CAT | K136 | - | - |
| 17 | | | | G-G | |
| 18 | | | | SG-GS | |
| 19 | | | | GP-PG | |
| 20 | | | | PG-GP | |
| 21 | | | | GPG-GPG | |
| 22 | | | | GP-PG | C450A |
| 23 | | | | | C450A/E409C |
| 24 | | | | | C450A/E409V |
| 25 | | | | | C450A/D432V |
| 26 | | | | | C450A/K488N |
| 27 | mRFP-LOV | mRFP | A225 | SG | C450A, EAAKGS |
| 28 | | | | | C450A, EAAKGS, E409V |
| 29 | | | | | C450A, EAAKGS, L493V |
| 30 | | | | | C450A, LAAKGS, E409V |
| 31 | | | | SSGSG | C450A, EAA |
| 32 | | | | | C450A, LAA |
| 33 | MCP-SoxS | MCP | N27 | GS-SG | C450A |
| 34 | | | N27 | | C450A, K488N |
| 35 | | | A56 | | C450A |
| 36 | | | A97 | | |
| 37 | | MCP-SoxS | linker in between | G-S | |
| 38 | AcrIIA5-E76-LOV | AcrIIA5 | E76 | - | C450A |
| 39 | | | | | C450A/K488N |
| 40 | SpyCas9-E566-LOV | SpyCas9 | E566 | SG-GS | - |
| 41 | | | | | C450A |
| 42 | | | | | C450A/D432V |
| 43 | | | | | C450A/P456L |
| 44 | | | | | C450A/D459E |
| 45 | | | | | C450A/K488N |
| 46 | SpyCas9-T1048-LOV | SpyCas9 | T1048 | SG-GS | - |
| 47 | | | | | C450A |
| 48 | | | | | C450A/P456L |
| 49 | | | | | C450A/D459E |
| 50 | | | | | C450A/D432V |
| 51 | | | | | C450A/K488N |
| 52 | SpyCas9-E566-cpGR2 | SpyCas9 | E566 | SG-GS | - |
| 53 | SpyCas9-T1048-cpGR2 | SpyCas9 | T1048 | SG-GS | - |
| 54 | dSpyCas9-E566-LOV-VPR | dSpyCas9 | E566 | SG-GS | - |
| 55 | | | | | C450A |
| 56 | | | | | C450A/K488N |

**Supplementary Table 4 | Genomic target sites of the sgRNAs used in this study.** Spacer sequences are marked in bold, PAM motifs are underlined.

| Cas protein | Target locus | Sequence 5’-3’ | Source |
|---|---|---|---|
| *Spy*Cas9 | *CCR5* | **TGACATCAATTATTATACAT**CGG | [8] |
| | *CFTR* | **AATGGTGCCAGGCATAATCC**AGG | [7] |
| | *AAVS1* | **GGGGCCACTAGGGACAGGAT**TGG | [11] |
| | TetO | **TCTCTATCACTGATAGGGAG**TGG | [10] |
| *Sau*Cas9 | *EMX1* | **GGCCTCCCCAAAGCCTGGCCA**GGGAGT | [6] |
| | *GRIN2B* | **GAGAGTAGGCTGGTAGATGGAG**TTGGGT | [6] |

**Supplementary Table 5 | Primers used for amplification of genomic loci.**

| Gene | Orientation | Primer sequence 5’-3’ |
|---|---|---|
| ***EMX1*** | Forward | GGGCCTGAGTCCGAGCAGAAG |
| | Reverse | CAAAAGGGAGATTGGAGACACG |
| ***GRIN2B*** | Forward | CAGGACGGCCAACACCAAC |
| | Reverse | GTGTATGCATACTCGCATGGC |
| ***AAVS1*** | Forward | GACAGCATGTTTGCTGCCTC |
| | Reverse | CTCCCTCCCAGGATCCTCTC |
| ***CCR5*** | Forward | GAAGGAAAAACAGGTCAGAG |
| | Reverse | CATTGCTTGGCCAAAAAGAGAG |
| ***CFTR*** | Forward | GACTAACCGATTGAATATGGAG |
| | Reverse | ATACACTTCTGCTTAGGATGA |
